# Late Fetal and Newborn Granulopoiesis but not Active Renin is Increased by Maternal Captopril Treatment During Perinatal Kidney Development

**DOI:** 10.1101/2022.05.15.491999

**Authors:** C Buckley, N L’Huillier, L Mullins, S Semprini, H Christian, JJ Mullins

## Abstract

Renin expression follows vascular development through the mouse kidney, regressing to glomerular poles by about P10, where renin is stored in dense core granules in juxtaglomerular cells. Homeostatic challenge to blood pressure causes release of active renin from the granules and recruitment of the renin lineage cells. We investigated the response to homeostatic challenge during late fetal development and following birth in a transgenic line expressing GFP under the renin promotor. Pregnant females were treated with water or captopril (30mg/kg/day), which inhibits angiotensin converting enzyme, from E15.5. We found an increase in renin transcription and expression by P1 following captopril treatment, with granulation increased at the glomerular poles and major arteries from E18.5. At P1, the granules showed a wide variation in electron density. Notably, rough endoplasmic reticulum was expanded in vascular smooth muscle cells (VSMCs) of captopril-treated pups at both time-points suggesting increased transcriptional activity. Paracrystalline material was observed in granules of captopril treated fetuses at E18.5 and in both treated and untreated pups at P1. Renin expression and some granules were confirmed in the kidney VSMCs by immuno-gold staining against GFP at E18.5. Importantly, we found no difference in active renin content between kidneys from treated and untreated pups at either age group. We therefore demonstrate a disconnect between granulation and active renin production in newborns when exposed to homeostatic challenge *in utero*.

## Introduction

The renin-angiotensin system (RAS) is one of the key enzymatic cascades regulating blood pressure, water and electrolyte homeostasis. In addition to its endocrine function, roles relating to the innate immune system in disease states such as inflammation ^1^, pre-B cell leukemia ^2^ and fibrosis, ^3,4^ tissue repair ^5^ or regeneration and trans-differentiation following glomerular damage ^6–9^ have been identified. The transcription and expression of RAS components in multiple utero-placental cell types has also been reported in humans and mice ^10^. Many of these extra-renal sites of expression may reflect local autocrine or paracrine actions of RAS.

Renin is an aspartyl protease, the function of which is to cleave its only known substrate, angiotensinogen, producing the decapeptide angiotensin (Ang) I by proteolytic cleavage at the N-terminus ^11^. Renin is synthesized as a pre-pro-enzyme of 50kDa in humans ^12^ and mice ^13^. In humans, the kidney is the only known organ to release active renin, whilst extra-renal sites secrete prorenin exclusively ^14^. However, some renin-producing tumours have been described ^15,16^. In addition, in the mouse, active renin has been found to be released by the submandibular gland ^17^ and by the adrenal gland ^18^.

Using antisera corresponding to the N-terminus, the C-terminus and the prosegment, Taugner *et al*. determined the fate of human prorenin during granule secretion ^19^. Both in humans and mice, prorenin can be released constitutively^20^, or packaged into low density protogranules that fuse and mature into dense core storage granules, as prorenin is converted to active renin (40kDa) for storage until controlled secretion is required ^21^. It is here that the 43 amino acid NH_2_ terminal pro-segment is cleaved off^20^, though the manner in which this occurs is controversial. Candidate prohormone convertases have been shown to cleave renin *in vitro*, however mouse knockout models for these enzymes did not alter active renin *in vivo*. For instance, whilst cathepsin B was shown to co-localise with renin inside dense core granules and showed site-specific cleavage of the pro-segment *in vitro*^22^, levels of active renin and prorenin remained the same in knockout mice as in controls^23,24^. There is evidence to suggest that after initial cleavage at the dibasic site, the pro-segment is slowly broken up until only the stable renin protein remains^25^.

In the adult kidney, active renin is released from a population of cells of the juxtaglomerular apparatus, the juxtaglomerular (JG) cells ^26^. JG cells are considered to be terminally differentiated from vascular smooth muscle cells (VSMC) since they possess VSMC characteristics and produce active renin. They derive from a pericyte lineage^27^; renin and pericyte markers such as αSMA and NG2 are coincidentally expressed during human kidney development, and primary cultures of pericytes isolated from human fetal kidney can be induced to express renin and form granules after stimulation with IBMX and forskolin^28^ Although some renin-expressing cells are present prior to vessel formation in the loose renal mesenchyme ^29^, renin expression in the developing mouse kidney essentially follows the formation of the arcuate and interlobar arteries ^30^ beginning on day E14 ^31^. As the renal artery divides further from interlobular arteries into arcuate arteries and afferent arterioles, renin expression follows this pattern and regresses in the larger arteries as the smaller arteries are formed, until it’s expression is confined to the classic juxtaglomerular location at approximately P10 ^31^. Sauter *et al* did not observe renin expression in the regions of greatest vascular growth and branching, suggesting that an established functional vessel wall is necessary for the onset of renin expression. However, the presence of renin during development suggests a physiological role of renin before blood pressure regulation commences.

In adults, when long term homeostasis is threatened, such as under chronic administration of angiotensin converting enzyme (ACE) inhibitors or low salt, vascular smooth muscle cells of the afferent arteriole, renal glomerular cells, mesangial cells and even interstitial cells are able to revert to a renin-producing phenotype^32^. This process, known as reversible metaplastic transformation^33^ or plasticity^34^, allows the production of renin in response to physiological challenges and is characterized by prorenin synthesis, granulopoiesis and the accumulation of mature granules to process prorenin into active renin. This pattern of recruited cells strongly resembles the ontogeny of renin development within the cortical vasculature^35^.

Post-natally, the presence of renin granules is unequivocal ^36,37^. However, few reports demonstrate the presence of renin granules in JG cells during renal development. In the pig, two reports showed the presence of granules in the mesonephros and metanephros (precursors of the mature kidney) ^37,38^. The presence of rare juxtamedullary juxtaglomerular renin granules was observed in Wistar rats at E18, with only occasional granules observed by E20 ^36^. Minuth *et al*., reported the presence of renal renin granules in NMRI mice ^39^, though transmission electron microscopy data was not shown. In human foetuses, a juxtaglomerular index (the number of granules/number of cells) of zero or below one was recorded ^40^ but granules were identified in metanephric tissue^41^.

It is, therefore, important to clarify whether renin dense core granules are present at the perinatal stage, and whether renin expression can be stimulated. We used transgenic animals expressing GFP under the renin promotor to show that ACE inhibition is a stimulus for renin transcription and perinatal granulopoiesis but is not a stimulus for storage of active renin.

## Experimental Procedures

Experiments were approved by the University of Edinburgh Animal Welfare and Ethical Review body (AWERB) and were conducted in accordance with the Animals (Scientific Procedures) Act 1986 and the Guiding Principles for Research Involving Animal.

### Transgenic animals and ACE inhibition studies

RenGFP^+/-^ mice^42^, on a C57Bl6/J background, were used throughout the study. Females were paired with males in the afternoon and were examined for vaginal plugs the following morning. The time of plug detection was termed E0.5. Pregnant mice were housed individually, and either water (control group; n=7) or captopril (30mg/kg/day in water; treatment group; n=5) was administered via gavage. Due to reports of fetal toxicity during maternal exposure in rats^43^, captopril (and water) gavage was started at E15.5. Male and female embryos were collected at E18.5 (n=5 water-treated mothers; n=2 captopril-treated mothers) and P1 (n=2 water-treated mothers; n=3 captopril-treated mothers) after CO2 administration, and both sexes were used in the studies performed.

#### Genotyping

Tails were collected and digested overnight following manufacturer’s instructions with DirectPCR lysis buffer (Viagen Biotech, 120-T). PCR was performed using VWR 2 mM MgCl_2_ Red Taq DNA polymerase 2X mastermix, following manufacturer’s instructions. Previously-published primers^44^ were used to determine the sex of the pups.

### Transmission Electron Microscopy

#### Ultrastructure Analysis

Embryonic and post-natal kidneys were cut in half and immersion fixed in 2.5% gluteraldehyde in 0.1M phosphate buffer (pH 7.2). Tissue was stored in this fixative for 4 hrs at room temperature, then transferred to a 10-fold dilution of buffer and stored at 4°C before preparation for electron microscopy by standard methods ^45^. Briefly, cells were post-fixed in osmium tetroxide (1% w/v in 0.1M phosphate buffer), then stained with uranyl acetate (2% w/v in distilled water), dehydrated through increasing concentrations of ethanol (70-100%) and embedded in Spurr resin (Agar Scientific). Semi-thin sections were cut and stained in toluidine blue for specimen orientation. Ultrathin sections (50-80 nm) were prepared using a Reichert Ultracut S Microtome, mounted on 200 mesh nickel grids, and stained lightly with uranyl acetate and lead citrate. Grids were viewed on a JEOL transmission electron microscope (JEM-1010, JEOL, Peabody, MA).

#### Immunogold labelling

Embryonic and post-natal kidneys were cut in half and immersion fixed in 3% paraformaldehyde in 0.1M phosphate buffer (pH 7.2) at room for 4 hours, transferred to a 10-fold dilution and stored at 4°C. Sections were prepared for immunogold electron microscopy by standard methods^46^. Briefly, segments were stained with uranyl acetate (2% w/v in distilled water), dehydrated through increasing concentrations of methanol (70-100%) and embedded in LR Gold (London Resin Company). Ultrathin sections (50-80nm) were prepared as above, incubated at room temperature for 2 hours with anti-GFP (Rabbit anti-GFP polyclonal Ab, Thermo Fischer Scientific, A11122, 1:1000) and for 1 hour with Protein A-15nm gold complex (British Biocell). All antisera were diluted in 0.1M phosphate buffer containing 0.1% egg albumin. As a control, the primary antibody was replaced by phosphate buffer/egg albumin. After immunolabelling, sections were lightly counterstained with lead citrate and uranyl acetate and were imaged with a JEOL transmission electron microscope as above.

#### Granule Electron Density Quantification

The mean pixel intensity was determined for each granule and normalized to the cytosol mean pixel intensity. Using this method the darker and more electron-dense the granule, the more negative the normalized mean pixel value.

### RNA extraction and quantitative real-time PCR analysis

Total RNA was isolated from frozen tissue using an RNeasy Micro Kit (Qiagen) according to manufacturer’s instructions. Kidney was homogenised in RLT buffer by shaking with a metallic homogenisation bead for 2 min at 30 Hz in a Retsch MM301 tissue disrupter (Haan). Genomic DNA was removed using a DNAFree Kit (Ambion) according to manufacturer’s instructions, and the RNA integrity verified using a Nanodrop and gel electrophoresis. cDNA synthesis was carried out using a High Capacity cDNA Reverse Transcription kit (Applied Biosciences) according to manufacturer’s instructions.

Mouse renin gene transcription was assessed by quantitative real-time PCR in a volume of 10 μl and monitored on a Roche Lightcycler 480 System using the Universal Probe Library. The primers and probe used for the assays were designed using the Roche Universal Probe Library Assay Design Centre and were obtained from Eurofins Genomic (EU). mRNA levels were normalised to 18S and HPRT mRNA. Mouse renin (UPL probe 16) forward primer: 5’-cccgacatttcctttgacc-3’, reverse primer: 5’-tgtgcacagcttgtctctcc-3’; 18S (UPL probe 77) forward primer: 5’-ctcaacacgggaaacctcac-3’, reverse primer: 5’-cgctccaccaactaagaacg-3’; HPRT (UPL probe 95) forward primer: 5’-cctcctcagaccgcttttt-3’, reverse primer: 5’ - aacctggttcatcatcgctaa-3’.

### Epifluorescence Microscopy

#### Imaging

Kidneys were dissected and placed in fresh phenol-free DMEM (10% FBS) on ice until imaging, where they were placed in a petri dish and visualized using a Leica MZ16 F stereomicroscope with top lighting onto a Hammamatsu Orca Flash 4 camera. The light source was a 100 W high-intensity mercury burner lamp, and a standard GFP emission filter was used (470/40nm).

#### Fluorescence intensity quantification

Images of kidneys were imported into FIJI, any region of interest (ROI) was marked and the mean fluorescence intensity measured. Background mean fluorescence intensity was measured, averaged and subtracted from the mean kidney fluorescence intensity. The area of the ROI was calculated, and the fluorescence intensity normalized to the area. All values are given in arbitrary fluorescence units (A.F.U.).

### Kidney renin activity

Kidneys were dissected from male e18 or P1 pups (collected from control or captopril-treated mothers; n = >5 per group), pooled, frozen and sent for analysis (Attoquant GmbH, Austria). Kidneys were powdered under liquid nitrogen and homogenized in phosphate-buffered saline (pH 7.4) by sonication. Protein concentration was determined by Bradford protein assay and normalized in all samples. The samples were diluted in an Ang I-stabilizing inhibitor buffer containing recombinant murine angiotensinogen (50 μg/ml). Samples were split in two parts and incubated at 37 °C in presence or absence of the renin inhibitor aliskiren. The incubation was stopped by acidification and the samples were extracted by Solid-Phase-Extraction prior to LC-MS/MS analysis. The difference of the Ang I level (in nmol/L) in both approaches allowed calculation of the renin-inhibitor-sensitive Ang I formation per hour per mg protein [((nmol/L Ang I) / h) / mg protein], which corresponds to the active renin activity in the tissue.

### Statistics

Data was analysed by Student t-test with Tukey’s post-hoc analysis and the level of significance was set to *P*<0.05. Error bars represent SEM with * p<0.05, ** p<0.01, *** p<0.001, **** p<0.0001 by two-way anova in conjunction with appropriate post hoc analysis.

## Results

Freshly isolated RenGFP^+/-^ kidneys were imaged using epifluorescence microscopy to visualise the extent of renin expression in fetuses and pups at E18.5 and P1 with and without captopril treatment of mothers (Fig. 1A). As expected, renin was expressed throughout the vasculature at both time points and expression at P1 appeared more extensive than at E18.5. By calculating the gross fluorescence intensity from the entire kidney, it was possible to show that whilst there was no difference between treatments in E18.5 kidneys, by P1 captopril treatment had significantly increased GFP expression, suggesting that renin expression was more extensive (Fig. 1B). This was verified using quantitative real time PCR (qRT-PCR) for renin, where renin transcription was only significantly increased in newborns from captopril-treated mothers (Fig. 1C).

**Figure 1.**
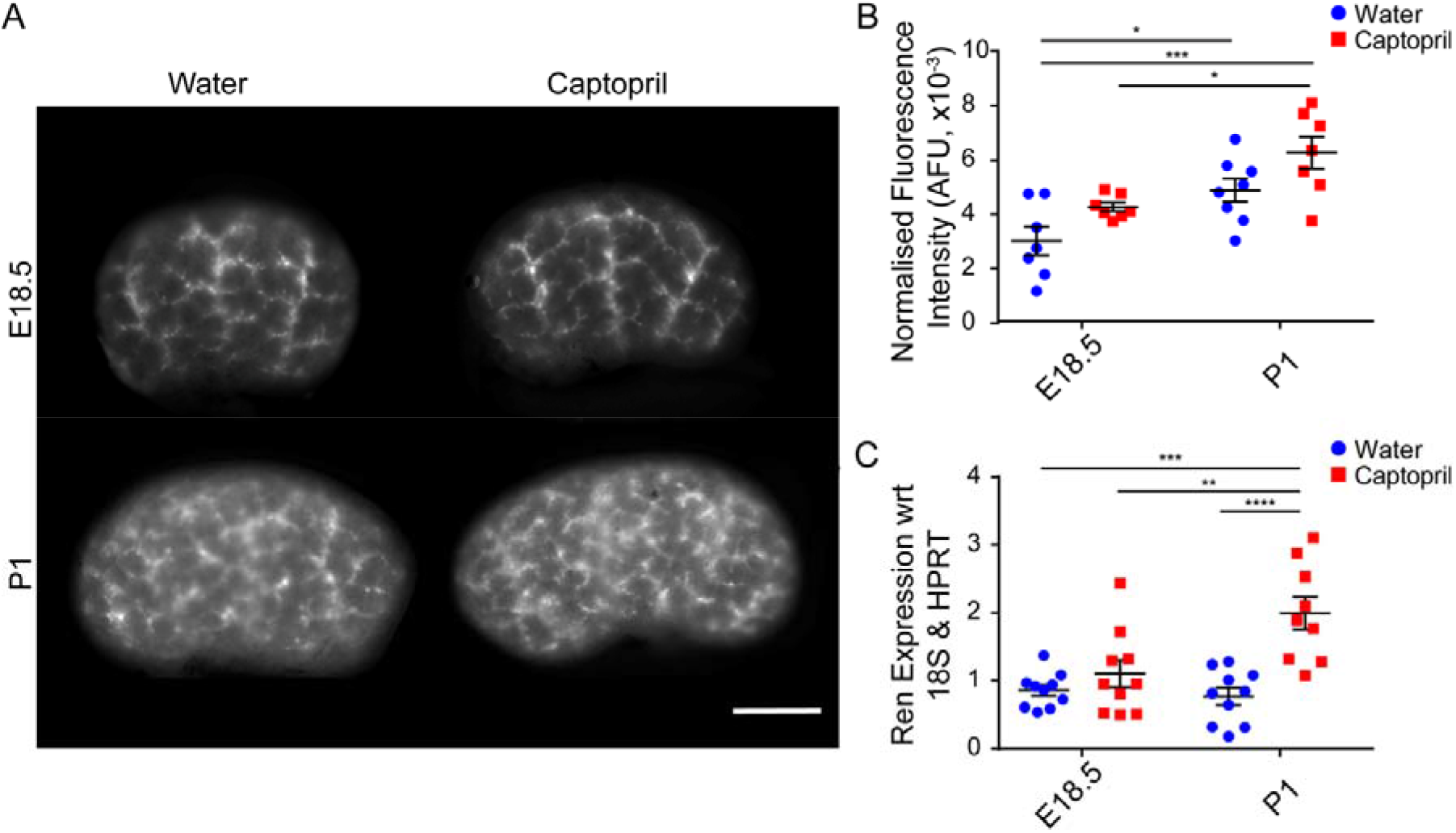
Fetal and post-natal renin expression after maternal water or captopril treatment. **A**. Kidneys from E18.5 and P1 RenGFP^+/-^ mice were dissected and imaged on an epifluorescence microscope to visualize renin expression patterns in controls and after treatment with captopril (30mg/kg/day captopril from E15.5). Scale bars - 0.5mm. **B**. Quantification of mean GFP fluorescence intensity in control (blue; E18.5 n=7, P1 n=8) and captopril-treated (red; E18.5 n=7, P1 n=7) RenGFP^+/-^ kidneys, normalized to the kidney area as imaged using epifluorescence microscopy. **C**. qRT-PCR quantification of renin transcript levels in control (blue; E18.5 n=10, P1 n=10) and captopril-treated (red; E18.5 n=10, P1 n=9) RenGFP^+/-^ kidneys. All error bars represent S.E.M, two way ANOVA with Sidak post hoc test performed for multiple comparisons, **:p<0.01, ***:p<0.001.

To determine whether renin expression was indicative of granulation within the renin-expressing cells, we performed electron microscopy ultrastructure analysis of the cells at the glomerular pole (Fig. 2i) and along major arteries (Fig. 2ii). Sparse granulation was seen in control E18.5 kidneys at the glomerular poles and along arteries, but granulation was considerably increased in captopril-treated, E18.5 kidneys (Fig. 2). More extensive granulation was observed at P1 (Fig. 3), both at the glomerular poles (Fig. 3i) and along major arteries (Fig. 3ii).

**Figure 2:**
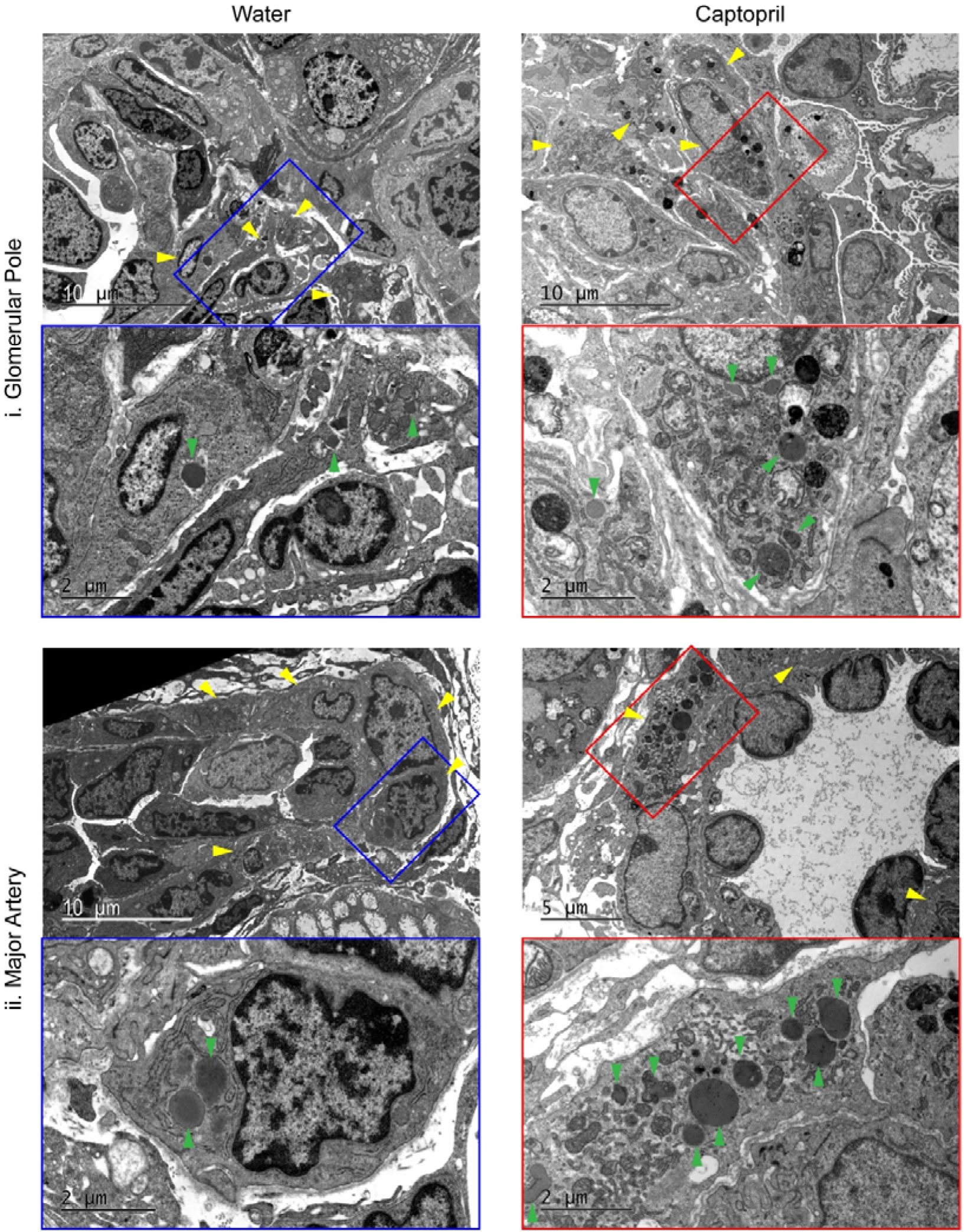
Electron microscopy ultrastructure analysis of E18.5 vasculature after maternal water or captopril treatment. Electron micrographs were taken of control (blue box) and maternal captopril-treated (30mg/kg/day captopril from E15.5) E18.5 embryos (red box) from i. the glomerular pole region and ii. major arteries, showing an overview of the region and a zoomed in renin-expressing cell. Yellow arrows indicate renin-expressing cells; green arrows indicate dense core granules. Scale bars represented individually.

**Figure 3:**
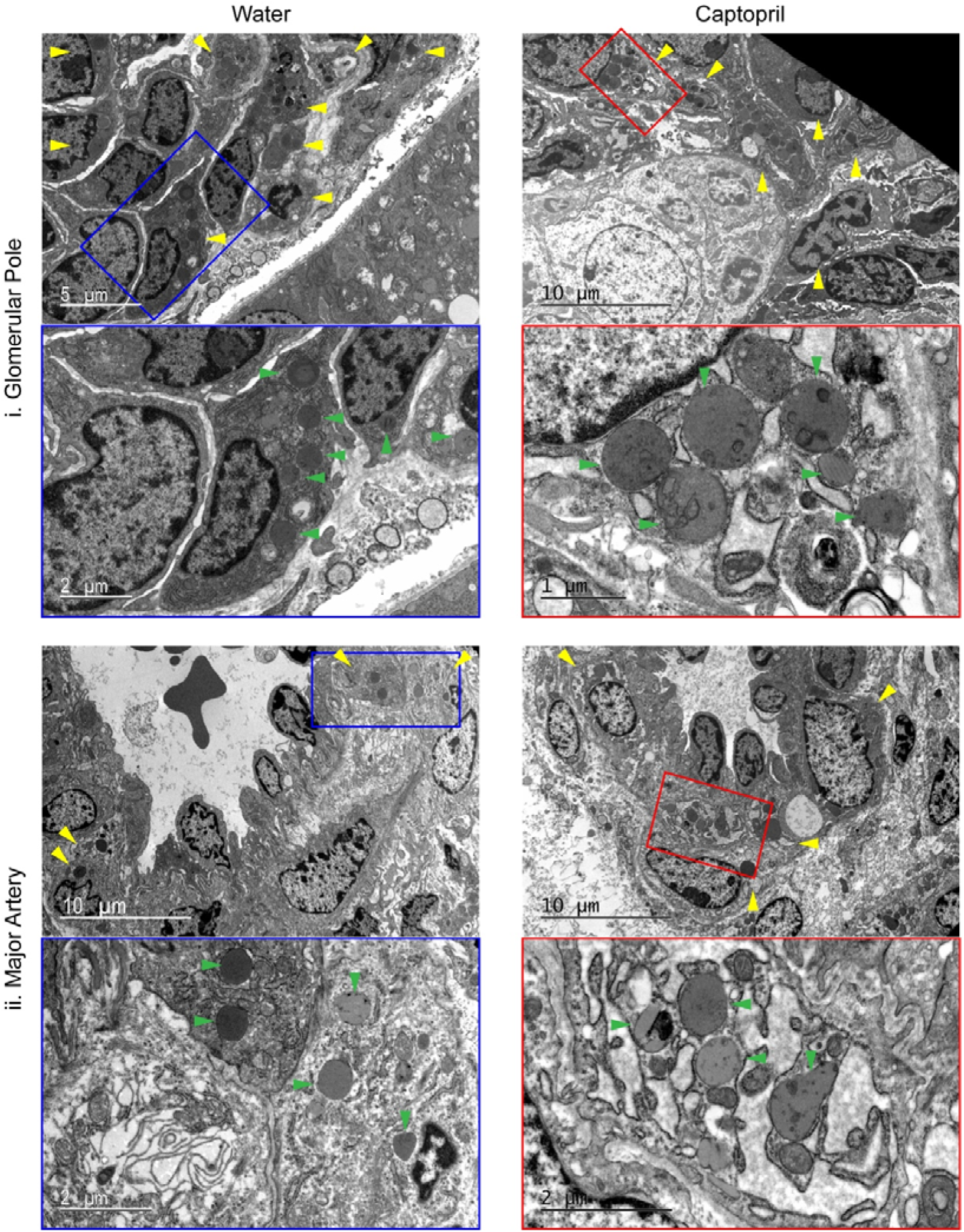
Electron microscopy ultrastructure analysis of P1 vasculature after maternal water or captopril treatment. Electron micrographs were taken of control (blue box) and maternal captopril-treated (30mg/kg/day captopril from E15.5) from i. the glomerular pole region and ii. major arteries, showing an overview of the region and a zoomed in renin-expressing cell. Yellow arrows indicate renin-expressing cells; green arrows indicate dense core granules. Scale bars represented individually.

The glomerular pole is the nominal location of juxtaglomerular cells, therefore granulated cells are necessarily renin-expressing. However, the vasculature only expresses renin during development and this expression pattern changes over time. To validate our method of identifying granulation in renin-expressing cells, we took advantage of the cytoplasmic GFP expression within and renin-expressing cells in the RenGFP^+/-^ mice by performing anti-GFP immunogold staining. To ensure that the immunogold stain is visible, images have been contrast-enhanced, and examples of the immunogold labelling highlighted using red arrows (Fig. 4). Immunogold staining can clearly be seen in the vascular smooth muscle cells (VSMCs) of major arteries under control conditions, but not in the underlying endothelial cells (Fig. 4A). Similarly, GFP^+^ staining is visible in VSMCs after captopril treatment (Fig. 4B).

**Figure 4:**
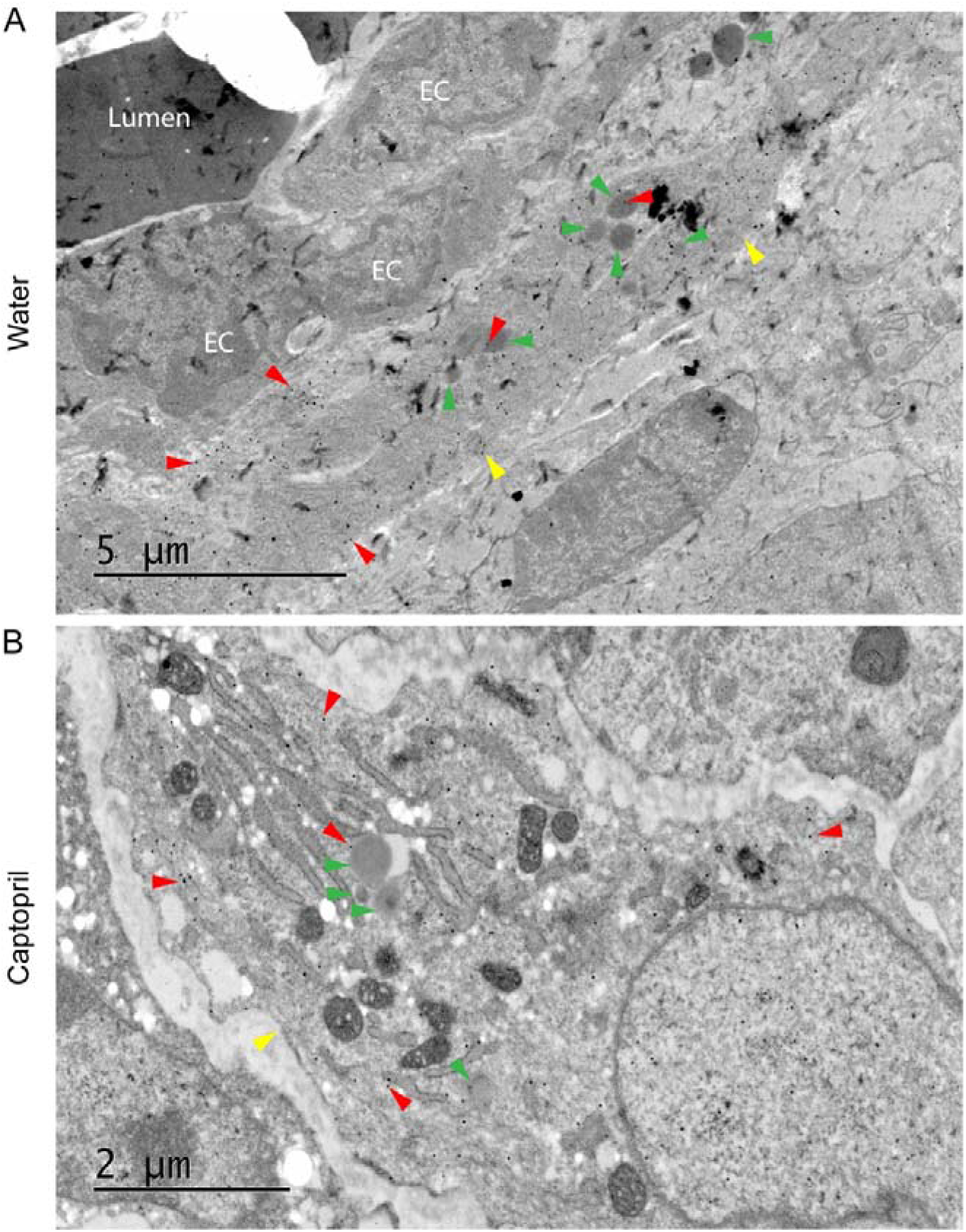
Electron microscopy of immunogold-labelled GFP verifies the presence of GFP-positive renin-expressing smooth muscle cells with granules. Electron micrographs were taken at E18.5 from **A**. control and **B**. maternal captopril-treated (30mg/kg/day captopril from E15.5) kidneys. Lumen = artery lumen; EC = Endothelial cell; Yellow arrows indicate renin-expressing cells; green arrows indicate dense core granules; red arrows indicate immunogold labelling examples. Scale bars represented individually.

Having validated our method of identifying renin-expressing cells, the ultrastructure of these cells was interrogated using electron microscopy. Granules within renin-expressing cells from captopril-treated E18.5 embryos appeared more electron dense compared to water-treated controls, while those in P1 samples exhibited a greater range of electron density after captopril treatment (Fig. 5A-B). To quantify this, the mean pixel intensity was determined for each granule and normalized to the background cytosol mean pixel intensity. The darker and more electron-dense the granule, the more negative the normalized mean pixel value. This confirmed the broadly equivalent electron density of granules from untreated E18.5 and P1 renin-expressing cells, however captopril-treated E18.5 renin-expressing cells showed a lower pixel intensity, indicating more electron dense granules. As expected, there was a more pronounced spread of electron densities of granules from captopril-treated P1 renin-expressing cells (Fig. 5C).

**Figure 5:**
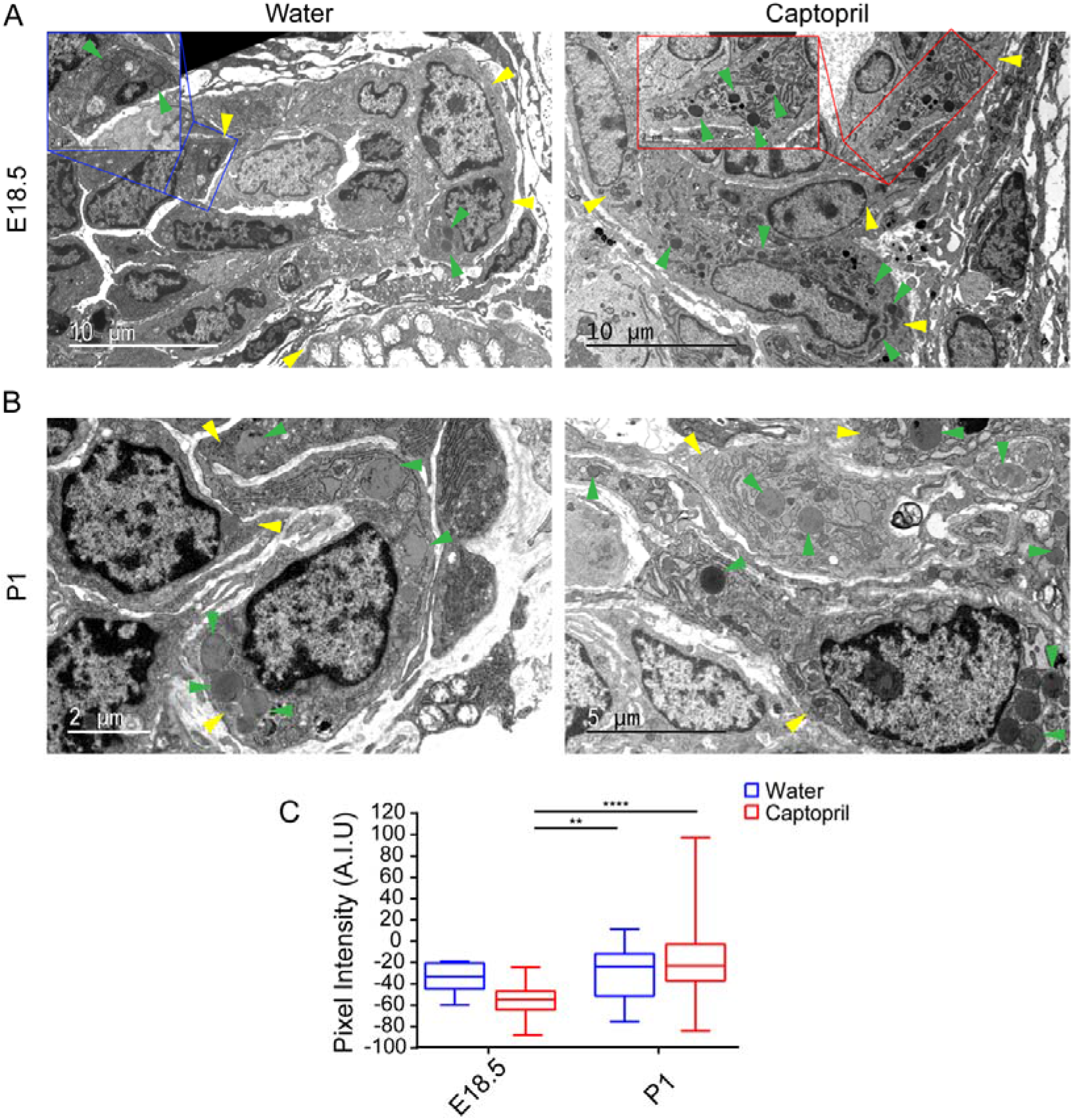
Electron microscopy of cellular ultrastructure reveals differences in granule electron densities between groups. Electron micrographs were prepared from **A**. E18.5 and **B**. P1 kidneys, of control (blue box) and after maternal captopril-treatment (30mg/kg/day captopril from E15.5; red box). **C**. Analysis of granule electron densities, where negative granules indicate a higher electron density. A.I.U = arbitrary intensity unity. Green arrowhead = dense core granule; yellow arrowhead = renin-expressing cell. Scale bars represented individually. Error bars represent outlier values in box and whisker plot, two way ANOVA with Sidak post hoc test performed for multiple comparisons, **:p<0.01, ***:p<0.0001.

Granules at all stages of granulogenesis were seen across all four groups. Of particular interest were protogranules containing paracrystalline material (Fig. 6A-B). These paracrystalline structures, when magnified sufficiently, were shown to contain regular lattices of proteins (dashed black arrows and black boxes; Fig. 6B-C). Often these also contained small vesicles within the membrane (cyan arrowheads). There were numerous instances of paracrystalline regular lattice material apparently lying outwith a granule membrane (Fig. 6C-D), again with pronounced vesicles in close proximity to the crystalline matter. The paracrystalline material was seen at E18.5 only after captopril treatment, and at P1 in both water and captopril-treated pups.

**Figure 6:**
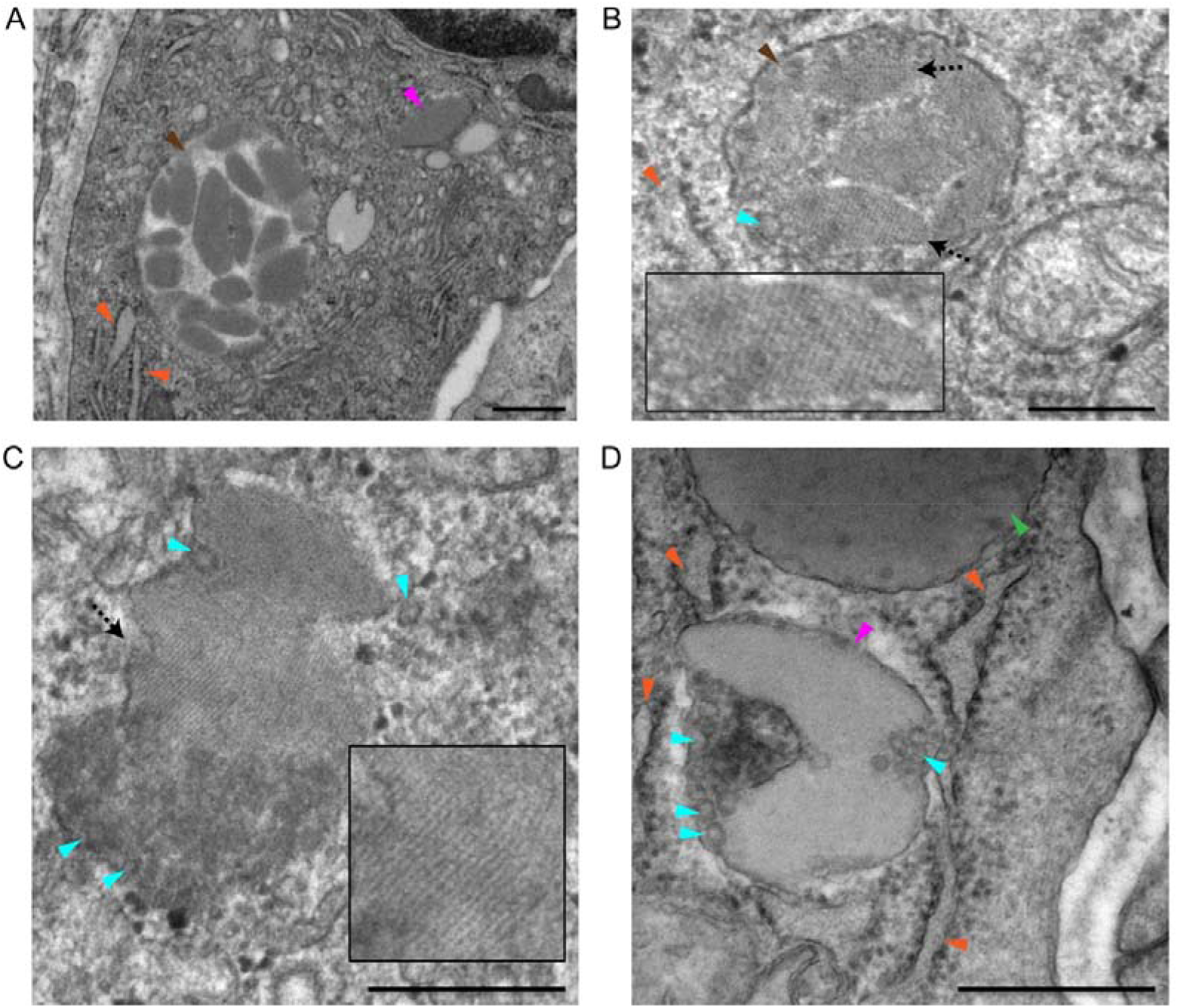
Electron microscopy of granular ultrastructure within renin-expressing cells. **A**-**B.** Electron micrographs of paracrystalline material accumulating within a granule-like membrane. **C** – **D.** Electron micrographs of paracrystalline material out-with an obvious granule membrane, with small vesicles closely located. Green arrowhead = granule; Cyan arrowhead = vesicle; orange arrowhead = endoplasmic reticulum; pink arrowhead = paracrystalline material out-with granule membrane; dashed arrow = paracrystalline material lies perpendicular to paracrystalline lattice; brown arrow = paracrystalline structures within membranes. Scale bars - 0.5 μm.

Another striking ultrastructural alteration brought about by maternal captopril treatment was the extent of rough endoplasmic reticulum (RER) dilation (Fig. 7), suggesting an increase in transcription levels within the cell. This was clearly visible at E18.5 compared to the water controls (Fig. 7A, yellow arrowheads) but was particularly prominent at the P1 timepoint where RER in the water control renin-expressing cells closely resembled that at E18.5, however the RER in captopril-treated cells were vastly dilated and took on an electron-lucent appearance (Fig. 7B).

**Figure 7:**
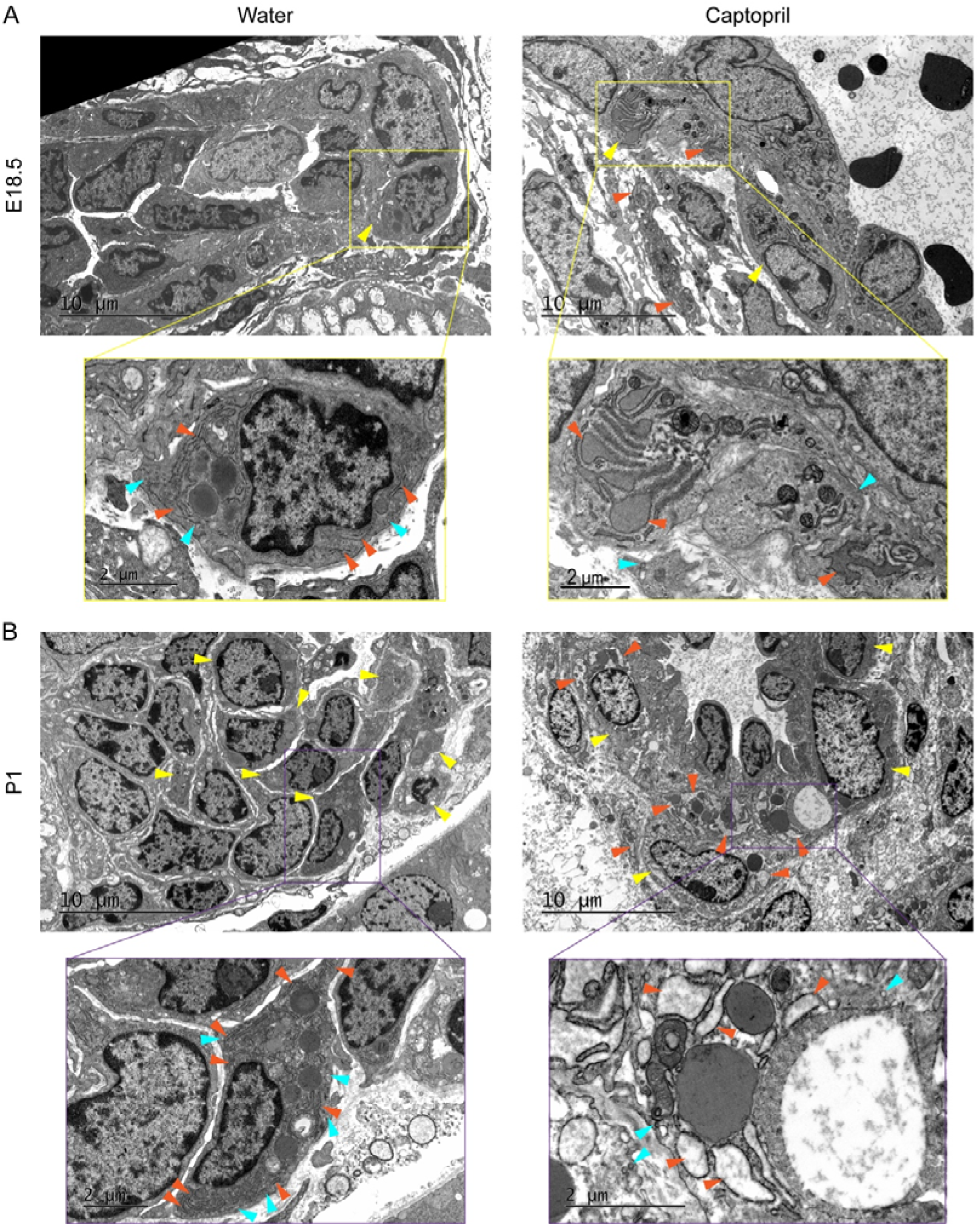
Electron microscopy of cellular ultrastructure reveals dilation of smooth muscle ER after captopril treatment. Electron micrographs were taken from **A**. E18.5 and **B**. P1 under control (water) and after maternal captopril-treated (30mg/kg/day captopril from E15.5) conditions. Yellow arrowhead = renin-expressing cell; cyan arrowhead = vesicle; orange arrowhead = endoplasmic reticulum. Scale bars represented individually.

The increase in renin expression and electron dense granules in P1 kidneys from captopril-treated mothers, coupled with the marked increase in RER size, would suggest that the levels of active renin may similarly be increased. To determine whether this was the case, pooled male kidneys from each timepoint (minimum 5 per group taken from mixed litters) were measured for renin activity (Attoquant GmbH). Results (Table 1) clearly show that the increased granulopoiesis seen in captopril-treated kidneys at P1 was not mirrored by a significant increase in renin activity.

**Table 1:** Renin activity measured in pooled e18 and newborn male kidney samples.

|  | **Renin Activity** |
| --- | --- |
|  | **[((nmol/L Ang I)/h)/ mg protein]** |
| **E18 Control** | 1404 |
| **E18 + Captopril** | 1247 |
| **P1 Control** | 2399 |
| **P1 + Captopril** | 2706 |
| **ADULT Control** | 3529 |

**Abbreviations:**

## Discussion

Previously it has been shown in the rat that granulation of renin lineage cells occurs only occasionally prenatally ^36^. Our preliminary studies suggested a complete lack of granulation in the mouse kidney at E16.5 (data not shown). We confirmed in the mouse the presence of renin immunogold-positive staining, together with occasional granulation, at E18.5, both at the glomerular pole region and within the kidney vasculature of control animals, with increasing granularity by P1. Most VSMCs contained few granules compared to adult recruited cells ^47^, confirming that the kidneys are still maturing after birth, and suggesting that the controlled secretion of active renin is not normally required perinatally. In the adult, pharmacological ACE inhibition up-regulates renin expression and production in recruited VSMCs as part of a feedback loop response. Using late gestation ACE inhibition, we observed an increase in dense core granules at E18.5 compared to controls, leading to a wide variation in granule electron density by P1. Renin transcript levels reflected dilated endoplasmic reticulum after captopril treatment in both E18.5 and P1 pups, but active stored renin did not show a parallel increase.

We used RenGFP transgenic mice to identify whether renin-expressing cells contained granules or not. This reporter strain had previously been shown to faithfully mark renin gene expression in the developing embryo, being detected in adrenals from E13 and kidney vasculature from E14. The reporter was also shown to regress towards JG cells with kidney development and exhibited recruitment of VSMCs on captopril treatment ^42^. We used late gestation (E15.5) captopril administration to mothers in this study, because previously in rats, captopril has been shown to decrease implantation numbers per litter and cause adverse effects on neonatal growth and survival^43^. Since captopril clearly crosses the placenta, we wanted to limit any toxicity.

It has been shown that only 12-15% of renin is packaged into granules in the rat around birth ^36^, the majority of renin presumably being constitutively secreted. The granules induced in mice by captopril treatment at E18.5 appeared to be classic dense core granules. Induction of granules in JG cells at E18.5 and P1 was also observed in preliminary studies following treatment of pregnant females with a 0.03%Na^+^, low salt diet (diet administered from E12.5; data not shown), showing that alternative stimulators of renin transcription also induced granulopoiesis. By P1, the granules were quite varied in electron density with increased numbers of low density or clear granules, which may reflect the lack of increased active renin being stored in dense core granules, suggesting that any increased prorenin expression led to constitutive secretion. Variation in granule electron density following renin gene stimulation in adult JG cells was also described when the human renin gene was used to rescue the mouse *Ren1d* knockout.^48^

Closer inspection of the proto-granules revealed paracrystalline material, often fringed by small vesicles and partially surrounded by membranes. Similar observations have been made on perinatal granules of the rat.**^36^** The appearance of these small vesicles and ‘ambiguous’ membranes^36^ around the granules in two separate species deserves further investigation, and may inform us about the early production of granules, crystalline lattices (thought to be prorenin arranged in regular arrays) and /or the activation of renin.

Renin lineage cells express not only renin but also aldose-keto-reductase, Akr1b7, which exhibits the same pattern of restriction to JG cells during late kidney development, and recruitment under physiological challenge^49^. Like renin, *Akr1b7* transcription is regulated by cAMP signaling ^50^. Interestingly, Akr1b7 is not co-localised with renin in granules, but is found in the endoplasmic reticulum^50^, where excessive dilatation was observed following captopril treatment, in this study and previously^48,51^. Taken together, the ultrastructural observations imply that captopril-induced increase in transcription and granulopoiesis can outstrip correct packaging into dense core granules both perinatally and in adults.

The central question is whether increasing granulation following captopril treatment of renin-expressing cells in the developing kidney involves the same mechanism as the increasing granulation seen with VSMC recruitment postnatally? Treatment of newborn rats with an angiotensin II type 1 receptor inhibitor, Losartan, for the first 12 days post birth was shown to arrest kidney vascular maturation, resulting in a reduced number of thickened afferent arterioles, together with fewer, smaller glomeruli^52^. This suggests that the RAS is playing a key role in kidney development. Using targeted knockout of vascular versus tubular renin, the defects in kidney development were shown to derive entirely from loss of vascular renin expression^53^.

A number of key signals have been identified as playing a crucial role in renin lineage cell recruitment, including RBPj ^47^, CBP and p300 ^54^ and microRNAs miR330 and miR125b-5p^55^. Conditional knockout of the transcription factor, RBPj, led to a significant decrease in both glomeruli and number of renin cells in the JGA, which was evident by 1 month of age. Following low salt + captopril treatment of adult mice, any recruited cells had very few granules compared to controls ^47^. When the histone acetyl transferases, CBP and p300, which are co-activators of the cAMP response element in the renin promotor, were conditionally knocked out ^54^ it was shown that the loss of renin-expression only became apparent once the kidney was matured at P30. The two microRNAs, miR330 and miR125b-5p, were found to have opposite effects on renin lineage cell recruitment, miR125-5p being down-regulated while miR330 was up-regulated^55^. Conditional deletion of Dicer, the enzyme which produces active microRNAs, again led to severe vascular and glomerular defects in the developing kidney^56^. Taken together, these studies suggest that there are subtle differences between the timings of stimulatory signals controlling renin expression in perinatal renin-expressing cells versus postnatal JG and recruited cells. On top of this, the presence of additional secretory triggers such as sympathetic nerve activity and establishment of the macula densa, contribute to postnatal blood pressure, water and electrolyte homeostasis ^57^.

In conclusion, our results add credence to the proposition that the response to captopril of renin-expressing cells in the prenatal and perinatal developing kidney is subtly different from that of the adult kidney. Captopril treatment increases granulopoiesis in cells that are already capable of producing prorenin (which is presumably constitutively secreted) but this apparently outstrips the ability of the ER machinery to correctly fill the dense core granules with active renin.

## Abbreviations

S.E.M: standard error on the mean
A.F.U: Arbitrary fluorescence units
A.I.U: Arbitrary intensity units
HPRT: Hypoxanthine guanine phosphoribosyl transferase
VSMCs: vascular smooth muscle cells
ECs: Endothelial cells
RER: Rough endoplasmic reticulum

## Acknowledgements

We wish to thank Professors Stewart Fleming and Jörg Peters for contributions to pilot studies and helpful discussions.

